# Obesity dysregulates feeding-evoked response dynamics in hypothalamic satiety neurons

**DOI:** 10.1101/2025.05.22.655553

**Authors:** Marta Porniece, Jessica Baker, Charlotte D. Ausfahl, Stephen X. Zhang, Mark L. Andermann

## Abstract

Melanocortin-4 receptor-expressing neurons in the paraventricular nucleus of the hypothalamus (PVH^MC4R^) integrate hunger-promoting and hunger-suppressing signals to regulate satiety. Food consumption-evoked responses in PVH^MC4R^ neurons increase gradually during meal consumption to promote satiety, and disrupting this process drives massive obesity. These critical satiety neurons are strongly affected by a high-fat diet, yet the impact on their functional properties remains unknown. We used fiber photometry to track PVH^MC4R^ neurons’ responses to the consumption of drops of milkshake in animals fed a chow diet or a high-fat diet (HFD), both after obesity was established and after its reversal. PVH^MC4R^ neurons in HFD-fed animals showed greater consumption-evoked responses than chow-fed animals at the early stages of meal consumption, and these responses did not increase further during the meal. HFD-fed animals also showed reduced licking vigor and motivation to consume Ensure. Switching HFD-fed obese animals to a normal chow diet (NCD) re-engaged the motivation to consume Ensure, partially restoring early-meal neural responses to a lower level, but did not restore the increase in consumption-evoked response magnitude across the meal. These findings highlight functional alterations in hypothalamic satiety-promoting neurons in obesity and provide insight into the pathological neural consequences of an obesogenic environment.

## Introduction

Neurons in the paraventricular nucleus of the hypothalamus (PVH) that express the melanocortin-4 receptor (PVH^MC4R^) are critical for regulating food intake and maintaining energy balance^1–4^. Recent research has advanced our understanding of how PVH^MC4R^ neurons process these signals under normal physiological conditions^5,6^. In lean animals, PVH^MC4R^ neuron activation is regulated by integration of inputs from two antagonistic upstream neuronal populations: 1) hunger-promoting Agouti-related peptide (AgRP) neurons, which selectively engage and inhibit PVH^MC4R^ neurons by releasing neuropeptide Y (NPY) that acts on NPY1 receptors^7–10^ (among other mechanisms^11–14^), and 2) the satiety-promoting pro-opiomelanocortin (POMC) neurons, which release α-MSH peptide from synaptic terminals onto MC4Rs in PVH^2,5,6,15,16^. This dynamic interplay is further modulated by circulating levels of metabolic hormones such as leptin and insulin, which reflect the body’s energy state and act on AgRP and POMC neurons to fine-tune the downstream response in PVH^MC4R^ neurons^17–19^. During a meal, increased α-MSH and reduced NPY release together elevate the intracellular second messenger cAMP in PVH^MC4R^ neurons to promote satiety ^5,6,20–22^. This mechanism ensures that caloric intake aligns with the body’s energy needs, preventing overeating and promoting energy balance. Notably, animals with AAV-mediated expression of the cAMP-degrading phosphodiesterase PDE4D3-Cat in PVH^MC4R^ neurons exhibit hyperphagia and rapid weight gain along with altered intrinsic excitability of PVH^MC4R^ neurons and impaired sensitivity to feeding-related excitatory inputs^2,5,23^.

The functionality of this finely tuned hypothalamic circuit becomes compromised in obesity^24–27^. For example, rare genetic variants that decrease α-MSH release by POMC neurons lead to early onset, severe, and rapid weight gain^27–32^. These conditions involving reduced stimulation of the MC4R are effectively treated by administering MC4R agonists, such as setmelanotide. These agonists bind to and activate MC4R, mimicking the natural signaling that would normally suppress appetite and promote energy expenditure^32–35^. In contrast, diet-induced obesity (DIO) may be associated with a saturation of MC4R signaling as well as other neural circuit dysfunction (see below). Accordingly, while setmelanotide treatment acutely improves multiple metabolic parameters in DIO, chronic setmelanotide-induced PVH^MC4R^ neuron activation is not effective in reducing food intake and body weight in DIO^36,37^.

DIO results from the excessive consumption of calorie-dense foods and has been linked to hypothalamic inflammation, gliosis, and other forms of hypothalamic injury^24^. DIO dysregulates feeding circuits by desensitizing AgRP and POMC neuron responses to food and altering excitability and neuropeptide signaling ^25,38,39^. In DIO, AgRP/NPY neurons show increased spontaneous firing due to altered intrinsic excitability^40,41^, while POMC neurons exhibit a decrease in spontaneous activity due to a hyperpolarized membrane potential^42^. Furthermore, in vivo fiber photometry recordings reveal obesity-driven reductions in intragastric nutrient- or hormone-induced modulation of AgRP neurons, which may either promote or reduce food intake (e.g., via desensitization of AgRP responses to intragastric infusion of fat or blunting of ghrelin-induced AgRP neuron activation, respectively)^25,43^. Obesity also attenuates the rapid responses of AgRP neurons to sensory food cues and food consumption^25,38,44,45^. Additionally, obesity blunts the responses of AgRP and POMC neurons to a variety of hormonal inputs that vary between fasted and fed states, such as ghrelin, CCK, leptin, and insulin^25,40,41,46,47^. These disruptions impair PVH^MC4R^ neuron sensitivity to upstream inputs and predict weaker elevations in cAMP during feeding and a weaker meal-related increase in PVH^MC4R^ excitation, thereby compromising the critical role of these neurons in energy balance, meal size, and weight regulation^5,37,47,48^. In the PVH, long-term HFD exposure induces the loss of MC4R protein abundance and mitochondrial content in PVH^MC4R^ neurons, even though the number of PVH^MC4R^ neurons remains the same^48,49^. This loss is accompanied by diminished a-MSH expression in the hypothalamic arcuate nucleus, further suggesting that exposure to dietary fat induces alterations in α-MSH-MC4R signaling^48^. In summary, excessive dietary fat consumption disrupts melanocortin signaling by impairing upstream inputs to PVH^MC4R^ neurons and how PVH^MC4R^ neurons integrate these inputs to regulate behavior.

Here, we investigate whether and how obesity-related disruptions in melanocortin signaling manifest in functional changes in PVH^MC4R^ neuron responses during feeding. We aimed to understand the maladaptive plasticity in PVH^MC4R^ neurons that may contribute to overeating in DIO. We employed fiber photometry to track the real-time activity of PVH^MC4R^ neurons during meal consumption in both lean and obese states, as well as following the transition from HFD back to a normal chow diet. Our results provide new insights into the plasticity of hypothalamic satiety mechanisms in response to changes in diet and highlight the potential for targeted interventions to restore energy balance in obesity.

## Results

### Feeding-related responses in PVH^MC4R^ neurons are elevated early in a meal in HFD-fed animals

To assess the state-dependent integration of satiety signals in PVH^MC4R^ neurons in lean and obese states (Fig. 1a), we provided animals with *ad libitum* normal chow diet (NCD) or calorie-dense high-fat diet (HFD, 60% of calories from fat) from 5 weeks of age. We then selectively expressed the calcium sensor GCaMP6s (AAV-Syn-Flex-GCaMP6s) in PVH^MC4R^ neurons in MC4R-Cre mice, and placed an optic fiber above the PVH for fiber photometry recordings (Fig. 1b). After recovery from surgery, the two cohorts of animals were fed limited, calorie-matched amounts of NCD or HFD daily (∼9.5 kcal/day; Fig. 1c, d). This food restriction motivated the animals to perform a simple operant tone-conditioned feeding task for 4-5 weeks.

**Figure 1.**
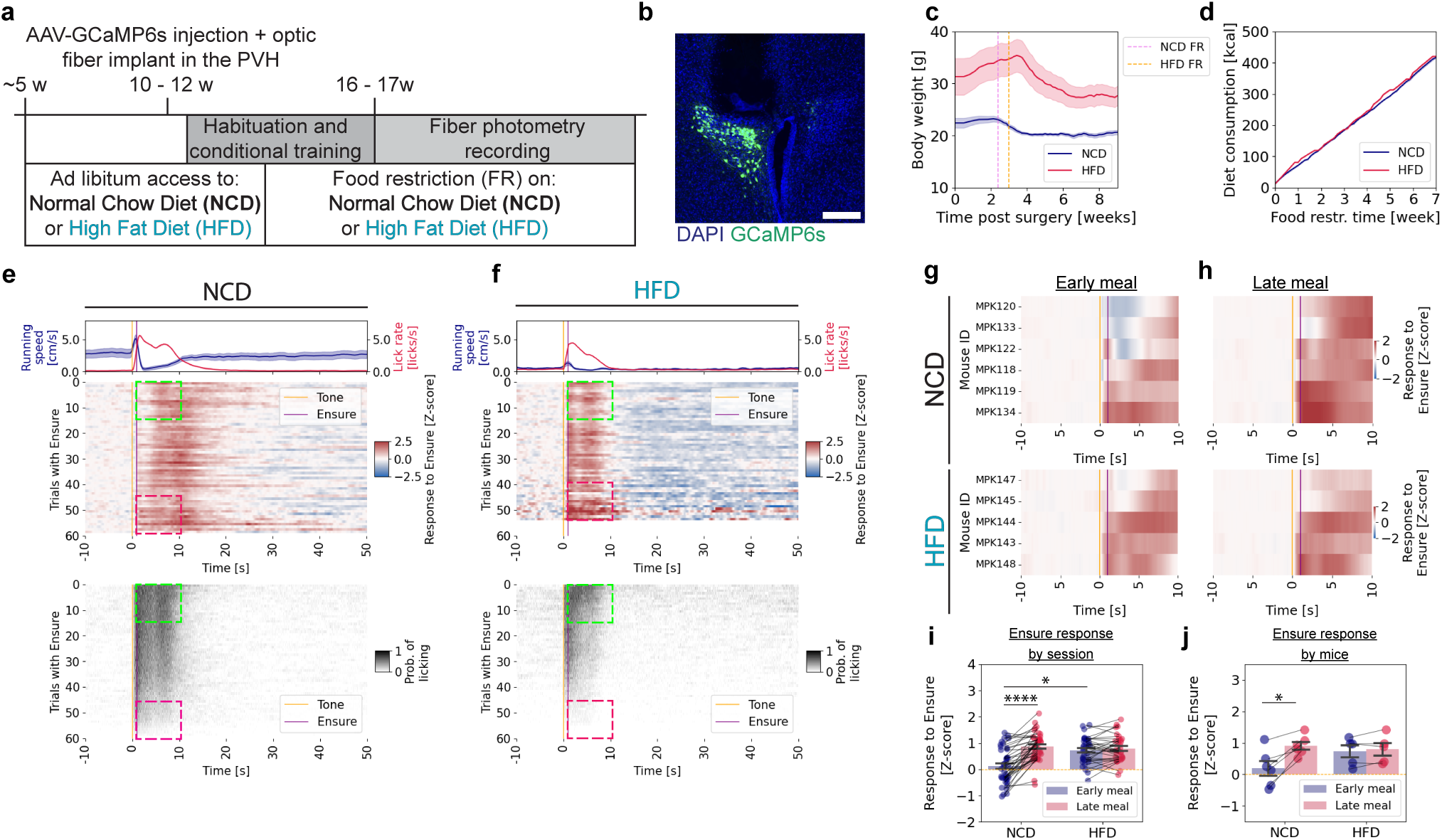
PVH^MC4R^ neuronal responses are elevated early in the meal in HFD mice. **a,** Schematic timeline of the experimental paradigm. **b**, Representative images showing viral expression of AAV-Syn-Flex-GCaMP6s and optical fiber placement in the PVH of MC4R^Cre/wt^ mice. Scale bar: 200 µm. **c**, Post-surgical bodyweight dynamics, showing initial ad libitum feeding and the onset of food restriction for each group (normal chow diet, food restricted: NCD FR; high fat diet, food restricted: HFD FR). NCD: 6 mice; HFD: 5 mice. **d**, Cumulative calorie consumption across both home cage food and Ensure during experimental recordings for NCD- and HFD-fed mice during food restriction. NCD: 6 mice; HFD: 5 mice. **e**, **f,** Mean heatmaps summarizing GCaMP6s photometry signals (top panel) and licking (bottom panel) from PVH^MC4R^ neurons of mice in NCD-fed (**e**) and HFD-fed (**f**) mice. Mean running speed and licking rates across all trials are shown above the heatmaps. Only recordings with a minimum of 30 successfully triggered trials were included. Trial structure: 10 s baseline followed by tone cue onset (t = 0 s) and Ensure delivery (t = 1 s) conditional on licking in the 1-s interval after tone onset. Early (first 15 trials) and late (last 15 trials) meal phases (quantified in **i, j)** are marked with green and magenta square insets, respectively. NCD: 39 recordings / 6 mice; HFD: 33 recordings / 5 mice. **g, h,** Mean heatmaps depicting trial-averaged PVH^MC4R^ neuron activity during early (**g**) and late (**h**) meal phases for individual NCD-fed (top panel) and HFD-fed (lower panel) mice by combining the sessions for each mouse. NCD: 6 mice, HFD: 5 mice. **i.** Z-scored PVH^MC4R^ neuronal responses during early and late meal phases for each recording in NCD-fed and HFD-fed mice. NCD: 39 recordings; HFD: 33 recordings. Linear mixed model (LME), p(NCD early vs. HFD early) < 0.001, p(Diet x Meal phase) < 0.001. **j**, Z-scored PVH^MC4R^ neuronal responses of each animal during early and late meal phases for all recordings in NCD-fed and HFD-fed mice. NCD: 6 mice, HFD: 5 mice. Paired, two-tailed test between early and late meal phases. **c – f, i – j**, Data are represented as the mean ± s.e.m. * - P <0.05, ** - P <0.01, **** - P <0.0001.

For both experimental groups, we monitored body weight and food intake in the home cage (Fig. 1c, d). During the fiber photometry recordings, we tracked Ensure consumption, licking vigor, and locomotion (Fig. 1e, f, Supplementary Fig. 1a, b). Upon food restriction, HFD-fed animals required a longer time to reach a steady-state level of reduced body weight (Supplementary Fig. 1c, d) and to engage in and learn the operant task (not shown). HFD-fed mice were ∼30% heavier than the NCD-fed mice, consistent with their elevated weight before the start of food restriction (Fig. 1c). Despite calorie matching to ensure comparable acute hunger states (Fig. 1d), the HFD animals consumed less Ensure volume during the experimental recordings (Supplementary Fig. 1a, b). Both NCD- and HFD-fed animals steadily reduced their licking rates from early to late phases of the session (Supplementary Fig. 1e, f). Compared to NCD-fed animals, food-seeking-related locomotion was negligible in HFD-fed animals (Fig. 1e, f top panels, Supplementary Fig. 1g), and it was further reduced to a minimum level late near the end of the meal (Supplementary Fig. 1g).

Using a head-fixed fiber photometry setup, we tracked calcium signals in real-time in parallel with licking behavior and locomotion (Fig. 1e, f, Supplementary Fig. 1h, i). Of all the trials in a session, we only analyzed the trials in which tone-evoked licking triggered Ensure delivery (“triggered trials”), as only these trials contributed to the gradual increase in satiety. NCD-fed animals consumed the palatable drops of Ensure from the beginning of the 60-minute meal, with licking vigor reducing throughout the meal as animals got satiated (Fig. 1e). Accordingly, after averaging all recordings (n = 39 sessions from 6 mice), we observed weak feeding-evoked responses early in the meal (first 15 trials) (Fig. 1e, g, h, Supplementary Fig. 1j). Similar to our prior study^5^, as the animals progressed through the meal, feeding-evoked responses gradually increased (Fig. 1e (top left panel)). This gradual increase in MC4R neuron activity and parallel reduction of licking vigor tracks the approach to the sated state as the animal continues to eat, which is likely to curb food intake once sufficient caloric intake is achieved^5^.

In contrast, HFD-fed animals (n = 33 recording sessions / 5 mice) stopped eating earlier during the experiment compared to the NCD-fed animals (Fig. 1e, f, lower panels; see also Supplementary Fig. 1g, h below). As discussed above, we matched levels of total daily calorie intake between NCD and HFD cohorts from the beginning of the food restriction and training phase of our protocol (to attempt to match acute hunger states across mice, Fig. 1c, d). Nevertheless, at the start of the meal, HFD-fed animals exhibited stronger feeding-evoked neural responses than NCD-fed animals, and these acute responses persisted until the end of the feeding bout (Supplementary Fig. 1j, k, left panels).

We next assessed the magnitude of PVH^MC4R^ neuron responses during the food consumption phase (0 – 10 s post-tone, restricted to trials with Ensure delivery). Specifically, we compared responses early in the meal (first 15 triggered trials) and late in the meal (last 15 triggered trials) in each recording. On average, these neural responses were minimal in NCD-fed animals in early trials (Z-score (NCD early) = 0.12 ± 0.11), but exhibited a robust increase in late trials (Fig. 1e (top panel); Fig. 1i) (Z-score (NCD late) = 0.86 ± 0.09), consistent with our prior study^5^. While some variation in early trial activity was observed among individual mice, all individual NCD-fed animals exhibited an overall increase in consumption-related neural responses in the late meal phase (Z-score (NCD early) = 0.19 ± 0.24, Z-score (NCD late) = 0.91 ± 0.13)) (Fig. 1g, h, j).

In contrast, PVH^MC4R^ neurons in HFD-fed animals were already highly driven during feeding in the early phase of the meal (Fig. 1f (top panel); Fig. 1g (lower panel), Fig. 1i) (Z-score (HFD early) = 0.72 ± 0.09), with no further increase observed throughout the meal (Z-score (HFD late) = 0.79 ± 0.09) (Fig. 1i). This pattern was consistent across all subjects (Z-score (HFD early) = 0.73 ± 0.19, Z-score (HFD late) = 0.80 ± 0.20) (Fig. 1h (lower panel), Fig 1j). Additionally, HFD-fed animals did not show a change in response from early to late in the meal when considering a longer window of 0-20 seconds post tone (Supplementary Fig. 1k). These experiments confirm a prior study^5^ showing that PVH^MC4R^ neurons are increasingly excited by food consumption as NCD-fed animals transition from hunger to satiety, and provide the first evidence that this within-meal change in neuronal responses is dysfunctional in HFD-fed animals. In particular, stronger-than-expected feeding-related responses in PVH^MC4R^ neurons at the start of a meal may explain the diminished motivation to consume Ensure in these obese animals (see Discussion).

### HFD alters neural and behavioral responses during the progression from hunger to satiety

We observed considerable inter-session variability in the number of trials with Ensure consumption required for the mice to become sated. Thus, we sought to subdivide the sessions by the number of trials required for satiation. In this way, we could compare neural and behavioral responses in the two groups of mice when matching for this index of motivation. However, because NCD-fed mice often consumed Ensure in all 60 trials in the experiments in the above experiments (Figure 1), we ran additional experiments in which we extended the session from 60 to 90 trials to allow for voluntary meal termination in both NCD-fed (n = 59 recordings / 7 mice) and HFD-fed (n = 54 recordings / 7 mice) animals.

During these extended recording sessions, lean animals continued to show progressive increases in feeding-related PVH^MC4R^ responses (Fig. 2a) throughout a larger number of trials than was possible in the earlier experiments. In these longer experiments, NCD-fed mice consumed Ensure during 70.1 ± 2.1 out of 90 possible trials, whereas HFD-fed animals only consumed Ensure during 52.9 ± 1.98 out of 90 trials (Fig. 2b). Consistent with our findings in Figure 1, in NCD-fed animals, excitatory responses increased over the meal, with a positive mean change in Z-score (ΔZ-score) between early and late in the meal of 0.52 ± 0.09. However, in HFD-fed animals, the increase was minimal (ΔZ-score = 0.12 ± 0.08; Fig. 2c). Similar results were observed across individual animals (ΔZ-score: NCD = 0.73 ± 0.07, HFD = 0.33 ± 0.10) (Fig. 2d). In these extended sessions, we first attempted to examine the across-trial changes in the animal’s motivation to consume food by measuring lick vigor during cue presentation. Throughout the meal, NCD-fed animals exhibited higher lick vigor than HFD-fed animals (Fig. 2e). In NCD-fed animals, lick vigor began to decline steadily after 50 trials, coinciding with the increase in PVH^MC4R^ responses as mice approached satiety. In contrast, in HFD-fed animals, lick vigor began to decline much earlier, after only 20 trials (Fig. 2e).

**Figure 2.**
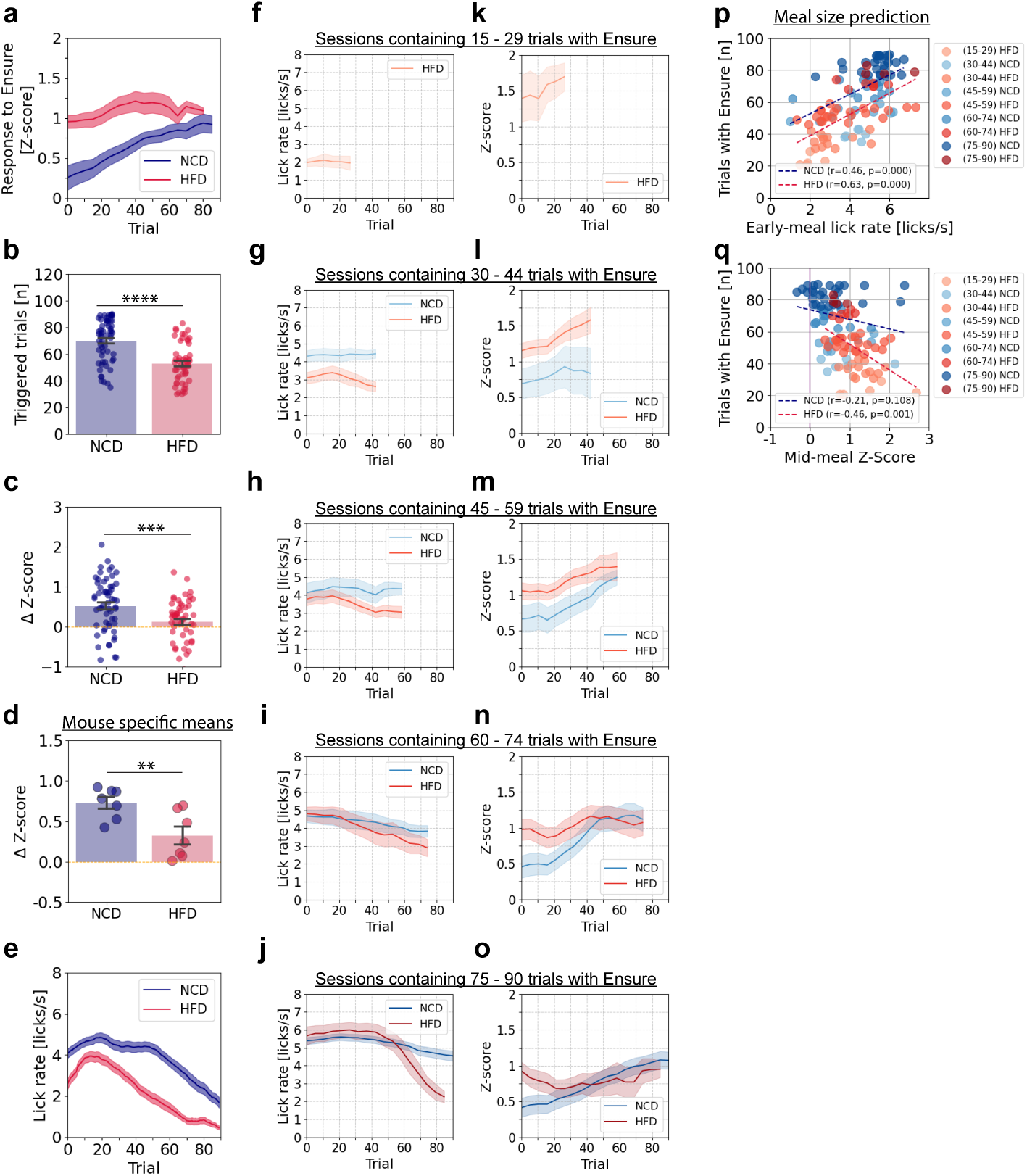
Mid-meal PVH^MC4R^ responses predict meal size in HFD mice. **a**, Single trial mean peri-licking PVH^MC4R^ neuron responses (1 – 9 s post cue) across 90 recorded trials from NCD-fed and HFD-fed groups (minimum 30 successfully Ensure triggered trials). **b**, Number of triggered trials during a 90-trial session for NCD-fed and HFD-fed groups. **c**, Difference in late-trial vs. early-trial neuronal responses (ΔZ-score) per recording for NCD-fed and HFD-fed groups. **d**, ΔZ-score averaged across sessions from individual NCD-fed and HFD-fed animals. **e**, Mean lick rate during audible cue presentation (t = 0 – 1 s) using the triggered trials as in **a**. **a – e,** NCD: 59 recordings / 7 mice, HFD: 54 recordings / 7 mice. **f – o**, Licking rate during the 1 s cue window (**f – j**), and single-trial average neuronal responses from the 1 – 9 s window post cue (peri-licking) (**k – o**) shown in each panel as the average across all sessions with meal size (i.e. # of triggered trials) within a certain range (15–29, 30–44, 45–59, 60–74, or 75–90 trials) in NCD-fed (blue shade) and HFD-fed (red shade) mice. Shaded error bars are ± s.e.m. across sessions with a given range of meal sizes. **p**, Correlation between meal size (number of triggered trials, i.e. trials with Ensure) and mid-meal neuronal responses (averaged across the 15^th^ to the 29^th^ triggered trial) for individual recordings, with trial range represented by light-to-dark shading (blue: NCD; red: HFD). Dashed lines indicate linear fits for NCD and HFD groups. **q**, Correlation between meal size (number of triggered trials) and mid-meal licking rate (averaged across the 15^th^ to the 29^th^ triggered trial) for individual recordings in NCD-fed and HFD-fed animals. Dashed lines indicate linear fits for NCD and HFD groups. **f – q,** Recordings for each meal size range: NCD (15–29): 0; HFD (15–29): 6; NCD (30–44): 6; HFD (30–44): 13; NCD (45–59): 10; HFD (45–59): 23; NCD (60–74): 13; HFD (60–74): 7; NCD (75–90): 28; HFD (75–90): 4. **b – d**, unpaired, two-tailed Student’s t-test. **p – q**, Pearson correlation. **a – o**, Data are represented as the mean ± s.e.m across sessions (**b, c, f – o**) or across mice (**a, d, e**). **- P <0.01, *** - P <0.001, **** - P <0.0001.

We hypothesized that on different recording days, animals’ hunger levels may have varied, potentially leading to different response magnitudes in PVH^MC4R^ neurons even when considering responses at the same time point in each session (Fig. 2b). Benefitting from the extended session duration, we could now stratify experiments by meal size before satiation (i.e. by the number of triggered trials where Ensure was delivered, in sessions with at least 15 triggered trials) and examine effects on the motivation to consume the food (licking vigor during cue) (Fig. 2f – j, Supplementary Fig. 2a) and the development of neural responses over phases of the meal (Fig. 2k – o, Supplementary Fig. 2b). First, we found that both HFD-fed and NCD-fed animals exhibited very variable lick rates early in the meal. Sessions with smaller meal sizes (fewer total triggered trials and lower caloric intake) exhibited lower lick vigor (i.e., the initial lick rate during the cue was 4-5.5 licks/s for NCD mice versus 2-5.7 licks/s for HFD mice) (Fig. 2f – j, p, Supplementary Fig. 2a).

We then analyzed the strengthening of PVH^MC4R^ responses throughout the meal across different meal sizes. In HFD-fed animals, sessions with short meals exhibited the highest neural responses early in the meal, and these responses reached their peak (Fig. 2k) as the lick rate fell to its minimum level (Fig. 2f), reflecting lower engagement and peak satiety. For short meal sessions with (30–44 triggered trials), HFD-fed animals exhibited elevated PVH^MC4R^ responses despite the reduced lick rate early in the meal. In longer meal sessions (45–59 and 60–74 triggered trials), licking vigor early in the meal was comparable between NCD- and HFD-fed groups, yet PVH^MC4R^ responses were initially larger and remained elevated in HFD-fed animals (Fig. 2g – h, l – m; Supplementary Fig. 2h – i). For the longest meal size (75 – 90 triggered trials), NCD-fed mice showed gradually increasing PVH^MC4R^ responses and consistently robust motivation to consume the food across the entire meal. In contrast, HFD-fed mice, despite demonstrating comparable lick vigor early in the meal, became satiated earlier, and their PVH^MC4R^ response magnitude did not increase throughout the session (Fig. 2I, j, n, o, Supplementary Fig. 2a – b). Overall, this analysis revealed that PVH^MC4R^ response magnitudes exhibited distinct dynamics throughout the meal in NCD-fed and HFD-fed animals, and also varied across sessions and mice within each group depending on meal size. Critically, despite both HFD- and NCD-fed mice having similar lick vigor early during a session (e.g., Fig. 2h), HFD-fed mice still showed elevated PVH^MC4R^ responses in the early trials (e.g., Fig. 2m). This indicates that the reduced motivation to consume food is likely not the primary driver of the elevated early session PVH^MC4R^ responses observed in HFD-fed animals. This finding supports the idea that obesity alters the underlying satiety-related signaling mechanism in PVH^MC4R^ neurons.

Further analyses revealed that the PVH^MC4R^ responses in HFD-fed animals negatively correlated with meal size (HFD: r = -0.46, p = 0.001) (Fig. 2q), whereas lick rate early in the meal positively correlated with the number of trials triggered in the session for both diet groups (Fig. 2p). This indicates that in HFD-fed animals, early PVH^MC4R^ responses and initial motivation to consume food early in a session predict subsequent meal size and caloric intake.

Similar to cue-evoked licking, licking during the reward period (1 – 9 s) post-tone in HFD-fed animals progressively declined with meal size (Supplementary Fig. 2c, d). We also assessed locomotion both before cue onset and after receiving the reward. Although HFD-fed animals exhibited reduced locomotion, this did not correlate with meal size (Supplementary Fig. 2e, f).

We next explored the relationship between within-meal increases in PVH^MC4R^ responses and overall meal engagement across sessions (Supplementary Fig. 2g). To this end, we correlated session-by-session differences in the number of triggered trials (a measure of meal size) with the magnitude of increase in PVH^MC4R^ response magnitude across the session. This analysis revealed a positive correlation in the NCD-fed group and a negative correlation in the HFD-fed group (NCD: r = -0.34, p = 0.012, HFD: r = 0.26, p = 0.048). This further highlights the decoupling of satiety circuit activity and meal size in HFD-fed mice. Interestingly, when we assessed sex-specific differences, we found that NCD-fed females exhibited slightly larger neural responses than males early in the meal – a trend that was even more pronounced in HFD-fed animals (Supplementary Fig. 2h).

Lastly, we also considered the possibility that the observed reduction in licking vigor and increased early-meal PVH^MC4R^ excitation in HFD-fed mice was due to insufficient caloric restriction in the context of the animals’ recent energy surplus. We therefore recorded sessions with overnight fasting in addition to the chronic food restriction protocol, to increase their hunger drive (Supplementary Fig. 2i – m). The additional fasting created an additional ∼9,5 kCal/day and ∼9 kCal/day calorie deficit for NCD- and HFD-fed mice, respectively. However, additional fasting did not significantly alter licking or PVH^MC4R^ response dynamics for either group (Supplementary Fig. 2i – m). Together, these results suggest that diet-induced obesity results in tonically high excitation of PVH^MC4R^ neurons that is not simply a consequence of caloric surplus but may reflect a disruption in the adaptive signaling required for proper meal termination and satiety regulation, potentially contributing to obesity maintenance.

### Effects of Diet Switching on MC4R Neuron Function and Feeding Behavior

To investigate whether PVH^MC4R^ neuron dysfunction in diet-induced obesity is reversible, we implemented a diet-switching protocol (Fig. 3a). A subset of animals previously fed a HFD (30 recordings / 7 animals) were switched to a NCD (42 recordings / 7 animals), while other NCD (52 recordings / 6 animals) and HFD (25 recordings / 4 animals) groups remained on their initial diets (NCD: 43 recordings / 6 animals; HFD: 23 recordings / 4 animals). This design enabled cross-comparisons among three groups: NCD-to-NCD, HFD-to-NCD, and HFD-to-HFD. Prior to this second phase of neural recordings (calorie-matched across groups), all animals were provided *ad libitum* access to their respective diets, followed by food restriction and re-habituation to operant conditioning.

**Figure 3.**
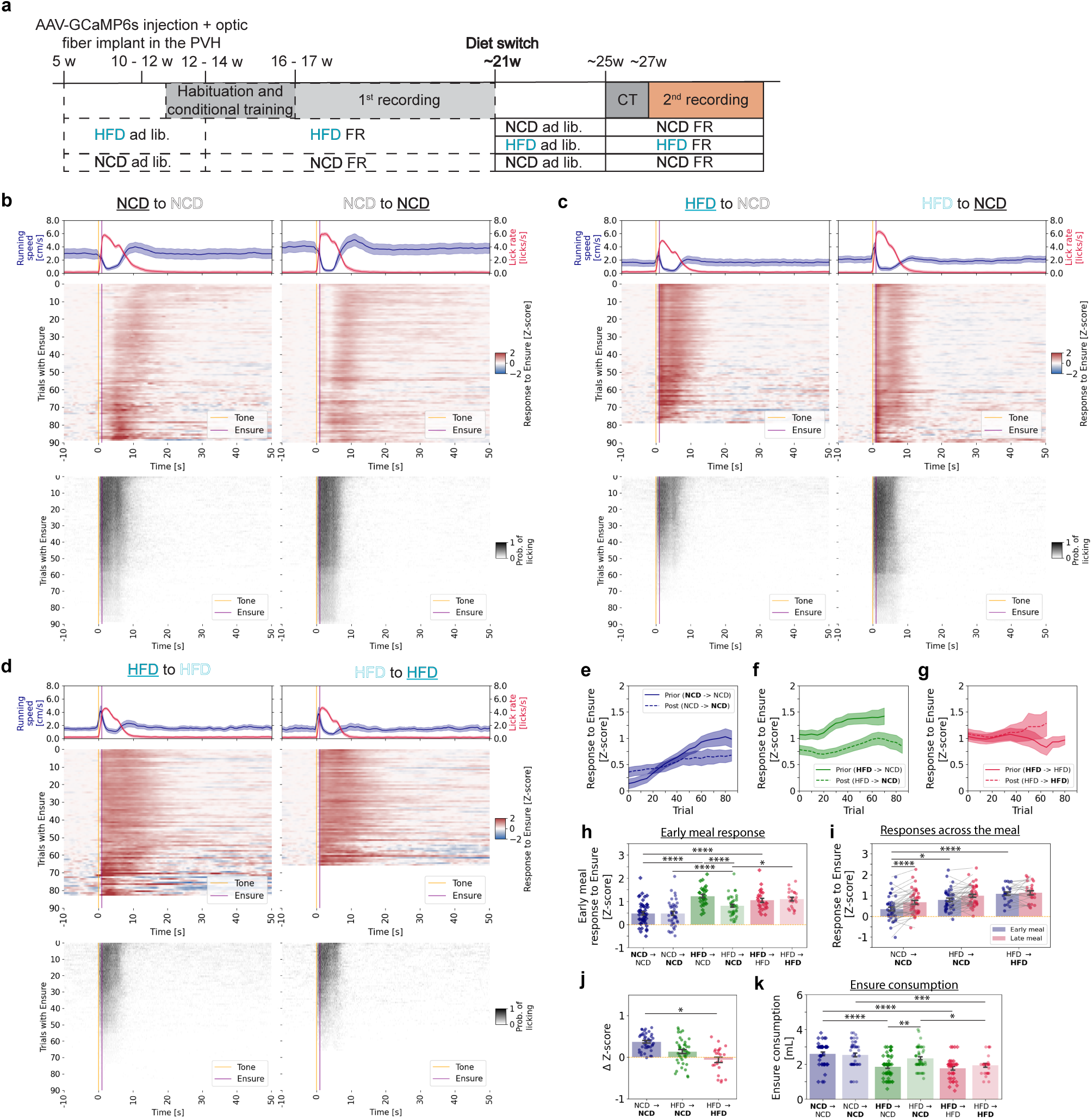
Partial reversal of PVH^MC4R^ neuron dysfunction and feeding behavior after diet-induced obesity. **a**, Schematic timeline of the experimental paradigm. Animals initially maintained on HFD were either switched to NCD (HFD to NCD) or remained on HFD (HFD to HFD) at ∼21 weeks of age. Control animals were maintained on NCD throughout (NCD to NCD). All groups underwent two recording sessions: one on their initial diet and another after diet switching (left and right panels, respectively, in **b – d**). **b**, Mean heatmaps summarizing GCaMP6s photometry signals (middle panel) and licking events (bottom panel) in the same NCD to NCD animals during the first phase of NCD diet (left, **NCD** to NCD) and second phase (right, NCD to **NCD**). Top panel: mean running speed and licking rate across all triggered trials. Only recordings with a minimum of 30 successfully triggered trials were included in the averaged heatmap analyses, while untriggered trials were excluded from the calculation of average photometry and behavior signals. **c**, Same as **b**, but for the HFD to NCD group of mice. **d**, Same as **b**, but for the HFD to HFD group of mice. **e – g**, Single-trial mean PVH^MC4R^ neuron responses during food consumption (1 – 9 s post cue) across triggered trials from all recordings in which at least 30/90 trials were triggered, for NCD to NCD (**e**), HFD to NCD (**f**), and HFD to HFD (**g**) groups, comparing pre-diet-switch (solid line) and post-diet-switch (dashed line) conditions. **h**, Z-scored PVH^MC4R^ responses during trials early in the meal (first 15 triggered trials) in NCD to NCD (blue), HFD to NCD (green), and HFD to HFD (red) groups, comparing pre-diet-switch (diamonds) and post-diet switch (circles) conditions. Each dot is a session. **i,** Z-scored PVH^MC4R^ responses post-diet switch during early vs. late parts of the meal (first 15 [blue] or last 15 trials [red]) for NCD to NCD, HFD to NCD, HFD to HFD post-diet switch. **j**, ΔZ-score for late vs. early trials aggregated for individual animals post-diet switch, categorized by groups of NCD to NCD, HFD to NCD, HFD to HFD. **k,** Ensure consumption during the individual photometry recording sessions, categorized by groups of NCD to NCD (blue), HFD to NCD (green), and HFD to HFD (red) groups, comparing pre-diet-switch (diamonds) and post-diet-switch (circles) conditions. **a – j,** Recording and animal numbers of the first diet exposure: **NCD** to NCD: 44 recordings / 6 mice; NCD to **NCD**: 43 recordings / 6 mice; **HFD** to NCD: 52 recordings / 7 mice; HFD to **NCD**: 42 recordings / 7 mice; **HFD** to HFD: 35 recordings / 4 mice); HFD to **HFD**: 23 recordings / 4 mice. **h, k,** Two-way Anova with Tukey post hoc test for before vs. after the diet switch within each condition and comparing first diet phase responses with each other and second diet phase responses with each other. **i,** Linear mixed model (LME), Value ∼ C(Condition, Treatment(’NCD | NCD’)) * Meal phase. **j**, one-way Anova, with Tukey post-hoc test. **b – k**, Data are represented as the mean ± s.e.m. either as error bars (**g – k**) or as a shaded area (**b – h**). * - P <0.05, ** - P <0.01, *** - P <0.001, **** - P <0.0001.

In this second phase of recordings, the NCD-to-NCD group again exhibited the previously observed PVH^MC4R^ neuron response potentiation across the meal, with the strongest transients occurring near the end of the 90-trial session (Fig. 3b, e, h). This activity correlated with vigorous licking during cue presentation throughout the experiment (Fig. 3b, e; Supplementary Fig. 3a). In contrast, during the initial phase of HFD consumption, the HFD-to-NCD group showed greater PVH^MC4R^ responses from the first bite but progressively reduced licking and ceased Ensure consumption by trial 80 (Fig. 3c, left panel, Supplementary Fig. 3c). After at least six weeks on NCD, these same animals displayed an increase in lick vigor during cue presentation (Fig. 3c, bottom right panel; Supplementary Fig. 3b), indicating a reinstatement of motivation to consume food. PVH^MC4R^ neuron responses remained attenuated relative to prior responses while on HFD for most of the experiment (Fig. 3f, h), but they did not show a noticeable increase in response magnitude across the meal, similar to the HFD-to-HFD group (Fig. 3i, Supplementary Fig. 3d). The HFD-to-HFD group maintained high early-meal PVH^MC4R^ responses (Fig. 3d, g, h) and ceased to engage in Ensure consumption even earlier in the session, never exceeding 60 trials. Licking vigor steadily declined across trials, reflecting a persistent drop in motivation to consume food (Supplementary Fig. 3c), as observed during their initial HFD exposure.

Further analysis revealed that only the NCD-to-NCD group exhibited a significant increase in PVH^MC4R^ responses from early to late meal stages (Fig. 3i). Interestingly, the HFD-to-NCD group displayed an intermediate pattern between the HFD-to-HFD and NCD-to-NCD, where there was a slight, but nonsignificant increase in PVH^MC4R^ responses across the meal (Fig. 3i; Supplementary Fig. 3d). When evaluating the PVH^MC4R^ response elevation across the meal (Fig. 3j), we confirmed a significantly smaller increase in response across trials in the HFD-to-HFD group (Fig. 3j). Notably, six weeks of NCD exposure in formerly HFD-fed animals failed to fully restore this range to NCD levels (ΔZ-score: NCD-to-NCD = 0.37 ± 0.03, HFD-to-NCD = 0.14 ± 0.05, HFD-to-HFD = -0.05 ± 0.07) (Fig. 3j). Despite this, the HFD-to-NCD group consumed a volume of Ensure comparable to the NCD-to-NCD group and significantly higher than the HFD- to-HFD group during individual recording sessions (Fig. 3k).

In summary, switching HFD-fed obese animals to NCD partially restored both neural activity (Fig. 3b, h, f) and feeding behaviors (Supplementary Fig. 3a). Specifically, in the HFD-to-NCD group, PVH^MC4R^ responses were closer to expected levels in the early meal stage (Fig. 3h) but did not show the same increase in response magnitude over the meal as the NCD-to-NCD group (Fig. 3i). This suggests a partial recovery of these neurons’ ability to integrate satiety signals (Fig. 3h). Moreover, the motivation to consume the Ensure, which had been significantly diminished in obese animals, was reinstated following the diet switch. Licking and food intake improved (Fig. 3k, Supplementary Fig. 3b), supporting the idea that a lean diet can rescue some aspects of both behavioral and neural dysfunction induced by obesity. These findings underscore the dynamic nature of the MC4R system in regulating feeding and energy balance. While diet-induced obesity impairs PVH^MC4R^ neuron function, transitioning to a healthier diet only partially restores the neurons’ ability to integrate hunger and satiety signals.

## Discussion

The present study sheds light on the role of PVH^MC4R^ neurons in integrating satiety signals during food consumption in lean and diet-induced obese mice, highlighting the alterations in neuronal activity and feeding behavior that occur with prolonged exposure to a high-fat diet (HFD). Our findings also confirm that in lean NCD-fed animals, PVH^MC4R^ neurons exhibit increased responses during feeding that are closely linked to meal progression and satiety^5^. Conversely, in HFD-fed animals, the response dynamics were impaired - PVH^MC4R^ neurons exhibit increased response magnitude early in a meal, consistent with the mice beginning the meal in a more sated state. This elevated response magnitude then stayed roughly constant throughout the meal (in contrast to the progressive increase in response magnitude in lean mice). This suggests a failure to properly integrate satiety signals as the meal progresses, or that a ceiling effect prevents a further rise in neural responses.

How do our findings in PVH^MC4R^ neurons relate to prior studies of DIO in upstream neural circuits involved in feeding behavior? Prior studies have shown that DIO mice exhibit increased spontaneous firing in AgRP/NPY neurons due to altered intrinsic excitability^40,41^. Moreover, these neurons are insensitive to leptin^41,50,51^. In contrast, POMC neurons exhibit a decrease in spontaneous activity due to a hyperpolarized membrane potential^42^. Moreover, long-term HFD exposure induces a loss of MC4R protein abundance in the PVH^MC4R^ neurons, even though the number of cells remains the same^48,49^. This loss is accompanied by diminished α-MSH expression in the arcuate nucleus, further supporting the notion that dietary fat exposure alters α-MSH-MC4R signaling^48^. The associated elevation in inhibitory NPY signaling (via Gi-coupled NPY1Rs) and reduction in excitatory α-MSH signaling (via Gs-coupled MC4Rs) would thus be expected to result in a decrease in cAMP in PVH^MC4R^ neurons at baseline (when the mouse is not consuming food). When we suppressed elevations in cAMP in PVH^MC4R^ neurons in a previous study using selective expression of a modified phosphodiesterase (PDE), mice developed massive obesity accompanied by a lower frequency of spontaneous activity and spontaneous postsynaptic potentials in PVH^MC4R^ neurons *in vitro*^5^. Thus, we initially predicted that PVH^MC4R^ neurons in HFD mice would be silent at baseline and more weakly activated during consumption, as seen in PDE-overexpression animals, but instead we observed the opposite effects in our experiments, where end-of-meal responses were similar to those in NCD mice.

We also considered other prior studies using *in vivo* fiber photometry recordings of calcium activity in DIO, which paint a more complex picture^25,43^. Acute delivery of various foods established that obesity attenuates the rapid sensory inhibition of AgRP neurons during food consumption^25,38,44,45^. Additionally, DIO blunts the responses of AgRP neurons to a variety of hormonal inputs, such as ghrelin, CCK, leptin, and insulin^25,40,41,46,47^. These disruptions likely prevent changes in cAMP in PVH^MC4R^ neurons in response to fast (consumption-related) and to more persistent (satiety-related) drops in AgRP neuron activity^25,37,45,48^. While these findings may help explain the lack of a gradual elevation in response magnitude over the meal in our HFD mice, they do not explain why response magnitude is *higher* in HFD mice at the start of the meal.

In lean animals, the gradual increase in PVH^MC4R^ neuron activity was accompanied by a reduction in licking vigor, a behavior closely linked to the development of satiety. The observed increase in consumption-evoked neural responses across trials was faster than in our previous study^5^, consistent with the fact that the animals satiated faster in the current study. This difference could be due to batch differences in body weight. In contrast, in HFD-fed animals, PVH^MC4R^ neurons showed unexpectedly strong responses early in the meal and remained similarly responsive throughout the meal. Given the higher body weight and metabolic demand in HFD animals, we would have expected them to engage in longer feeding bouts. Surprisingly, despite being food-restricted and calorie-matched to NCD-fed mice, HFD-fed mice displayed premature satiation. Further, the degree of this premature satiation was predictable from neural response magnitudes early in the meal.

Interestingly, the results from the diet-switching experiment provide insight into the potential reversibility of these impairments, as previous studies show recovery of expression of MC4R and α-MSH after returning to a low-fat diet for four weeks^48^. While the HFD-to-NCD animals in our study exhibited some recovery in motivation to consume food, as evidenced by an increase in licking vigor and larger meal size, the PVH^MC4R^ neuronal responses did not fully return to the levels observed in the NCD-to-NCD group. Notably, some drawbacks must be considered when interpreting the results of the diet-switching experiments in Figure 3. The prolonged expression of the calcium indicator and potential overtraining of the animals to perform the task may have contributed to the slightly less pronounced PVH^MC4R^ neuron response potentiation observed during the second recording after the diet switch to NCD. This is reflected in the lower ΔZ-scores compared to those in Fig. 2c.

Additionally, HFD-fed mice devalue NCD as a food resource, consistent with blunted AgRP neuron sensitivity to NCD and reduced intake of NCD^25,38^. A two-week withdrawal from long-term HFD exposure is insufficient to restore the AgRP neuron responses to sensory food detection and consumption^38^. In our study, although we returned mice to NCD for over 6 weeks before recordings, only a partial recovery of responses was observed. These findings suggest that the neuronal dysfunction in HFD-fed animals may not be entirely reversible with short-term diet intervention, indicating a potential long-term impact of HFD exposure on the plasticity of the PVH^MC4R^ neurons.

Although DIO increases the spontaneous firing of AgRP neurons^39,51^ and inhibits POMC neurons^42^, and AgRP neurons become less sensitive to leptin^39^, administration of a leptin antagonist in DIO mice promotes additional food intake^52^, Hence, our finding of elevated food consumption response early in the meal may be due to the persistently high level of leptin in DIO mice, both before and after the meal. In both cases, leptin is likely to saturate its receptor, leading to consistently elevated excitability of POMC neurons and reduced excitability of AgRP neurons. Under these conditions, cAMP levels in PVH^MC4R^ neurons may not return to a low level in between meals, potentially preventing a downward resetting of the strength of consumption-evoked synaptic inputs to PVH^MC4R^ neurons. Concurrently, we observed that calorie-restricted HFD mice, despite being calorie-matched to the NCD group and theoretically having the same acute caloric deficit while exhibiting increased adiposity and leptin levels, had small meal sizes during the recordings, as expected based on their higher early-session PVH^MC4R^ neuron responses.

This hypothesis is consistent with the counterintuitive finding from Sherrer and colleagues showing that clamping leptin at a *lower* level in DIO mice actually reduces body weight, despite the anorexigenic actions of leptin in healthy mice^53^. This may allow consumption-evoked responses to drop to a lower magnitude early in the meal, and the associated restoration in leptin sensitivity may also be involved in restoring the normal increase in response magnitude throughout the meal.

Our findings, together with the aforementioned effects of leptin antagonist administration in DIO mice, suggest that the mechanisms responsible for suppressing homeostatic feeding, such as those mediated by leptin, are at least partially intact in HFD mice and are engaged from the beginning of the meal. This indicates that the physiological “brakes” that typically signal satiety and reduce the drive to eat are functional to some extent. In addition, the persistence of feeding despite these satiety-promoting signals could also imply that a second, parallel motivational system may be at play. Specifically, circuits involved in processing the hedonic or rewarding properties of high-fat diet (HFD) food might be overriding or competing with homeostatic signals^54^. These reward-related pathways could be contributing to the decision to initiate and sustain eating, independent of true caloric need, particularly in the context of energy-dense, palatable food.

Future experiments, such as assessment of peptide transmission from inputs onto PVH^MC4R^ neurons in DIO, will be necessary to further investigate the mechanisms underlying these changes and to better guide therapeutic interventions.

## Methods

### Animal care

All animal procedures were approved and completed in compliance with the Institutional Animal Care and Use Committee at Beth Israel Deaconess Medical Center (BIDMC). All mice were single-housed in Innocage® plastic ventilated cages (Innovive) and kept in a room on a 12h light/12h dark cycle. Room temperature (18-22°C) and humidity were controlled within the rodent housing room. From five weeks of age to approximately 12 weeks of age mice were kept on an *ad libitum* diet of either standard mouse chow diet (LabDiet®, Formulab Diet Irradiated, 5008) or blue dye high-fat diet (Research Diets, D12492i, Rodent Diet) with 60 % of total kcal from fat, 20% from carbohydrates, 20%from protein. Both male and female MC4R-Cre^tg/wt^ (B6.129S4-Mc4rtm1Lowl/J) mice were included in the experiments. The number of mice used for each experiment is shown in each Figure. Experiments were not conducted in an experimenter-blinded model, but animals were randomly assigned to either a lean diet or a high-fat diet group.

### Chronic food restriction

After recovering from stereotaxic surgery, the mice were chronically food restricted either on normal chow pellets (Bio-Serv, Product # F0171, Dustless Precision Pellets, 500 mg, Rodent Grain-Based Diet, Lot # 295488.00) or blue dye high-fat diet for the remainder of the first round of photometry recordings (Figure 1 – 2). For the second recording phase (Figure 3), we food-restricted the animals on the same or on a different diet. Mouse body weight and food intake were recorded to keep both groups of mice at ∼80% of their original body weight (Supplementary Fig. 1c, d). Some recordings (Supplementary Fig. 2c – e) involved acute overnight fasting of the food-restricted animals the day before (16h), to increase motivation to eat. After the end of a recording that was preceded by acute fasting, the animal was refed with about twice the amount of food given during the standard food-restriction protocol to maintain body weight at ∼80% of the original body weight.

### Stereotaxic surgery

We performed stereotaxic viral injections and headpost implantation following a protocol that was described previously^5^. Briefly, AAV1-Syn-FLEX-GCaMP6s (Addgene, 100845-AAV1) was injected unilaterally into the PVH of MC4R-Cre^tg/wt^ mice (150 nL, Bregma: AP -0.6 mm, ML - 0.2 mm, DV -4.75 mm) at a titer of 1.05 x 10^13^ gc/ml. Single optic fibers with a metal ferrule (MFP_400/430/LWMJ-0.57_1m_FC-ZF1.25 (F)_LAF; Doric Fibers) were implanted above the PVH (Bregma: AP -0.6 mm, ML -0.25 mm, DV -4.6 mm) in each mouse. C&B Metabond (Parkell) was used to cement the titanium metal headposts onto the cranium following surgery. Mice recovered for 2 weeks post-surgery before any other intervention. Viral expression and accurate fiber positions were verified using the *post-hoc* histology.

### Behavioral training on wheels

Two weeks after surgery, animals started a 3-4 week protocol of habituation followed by behavioral training. Head-fixed mice were placed on a three-dimensional (3D)-printed standalone circular treadmill for three days at increasing intervals of 20, 30, and 45 minutes, respectively. On Days 4 through 8 (D4-8), animals were introduced to Ensure (Ensure® Plus nutrition shake, 1.47 kcal/mL) via manual administration through a syringe three times during the one-hour training session. Starting at D9, mice were head-fixed and placed on the circular treadmill with access to a lickspout for 25 minutes. Ensure was manually dispensed from the lickspout, and the mice were considered acclimatized when they licked regularly from this lickspout. Training then continued on the circular treadmill for another 30 minutes. Mice were further acclimated to the recording set-up by letting them freely run on the treadmill and self-administer Ensure from the lickspout. Animals then underwent classical conditioning to associate an audible cue with the delivery of Ensure, followed by operant conditioning in which mice would lick to trigger the Ensure delivery via a solenoid pump after an audible cue. Milkshake delivery speed was controlled by gravity and regulated by a solenoid pulse (5 x 150 ms pulses, 150 ms between the pulses, 3 μL per pulse, ∼25 μL total per trial) during all training sessions. For unconditional training, usually, 3 – 4 45-minute-long sessions with 75 Ensure deliveries after a cue (released at random intervals between 45 s and 70 s) were carried out per animal. For operant conditioning training, at least 3 training sessions were carried out to ensure that animals were robustly triggering the Ensure release after each audible cue. Operant conditioning training sessions were initially short and were then gradually extended across days as the behavior became more reliable. If animals (especially HFD-fed mice) were not progressing well through initial learning of this task, they underwent additional classical conditioning training sessions, and then restarted the operant conditioning.

### Head-fixation and food delivery

Mice ran on a circular treadmill with running speed measured by an IR beam while their head were fixed. Head-fixation reduces motion artifacts while recording neural activity. In all recordings, Ensure was delivered through a lickspout, with the following protocol: 10 x 150 ms pulses, 150 ms between the pulses, 3 μL per pulse, ∼50 μL total per trial. Milkshake delivery during recording was controlled using the Nanosec (https://github.com/xzhang03/NidaqGUI) photometry-behavioral system. The total consumed Ensure throughout the experiment was noted.

### Head-fixed photometry

Head-fixed photometry experiments were conducted as described previously^5,23^. For GCaMP6s recordings, excitation light from 465 nm (PlexBright LED 465nm, Plexon) and 405 nm (PlexBright LED 405nm, Plexon) LEDs were modulated by LED drivers and combined in a three-port mini-cube (FMC4_IE(400-410)_E(460-490)_F(500-550)_S, Doric) to transmit to the implanted optical fiber via a patch cord (NA 0.57, MFP_400/430/LWMJ-0.57_1m_FC-ZF1.25(F)_LAF, Doric). The emitted light was collected in a femtowatt photoreceiver (2151, Newport). Photometry excitation light was controlled by the Nanosec photometry-behavioral system^5,23^. Briefly, excitation light was modulated as interleaved pulses (465 LED ON/405nm LED OFF for 6 ms; 465 LED OFF/405nm LED ON for 14 ms). The respective datapoints of the 6ms pulse were extracted to quantify the photometry signal. The recording was conducted in the dark. Three to five recordings per mouse were performed in two conditions: 1) food restriction (FR) and 2) food restriction plus acute overnight fasting of FR mice prior to the experimental recording. Acute overnight fasting (16 h) was implemented to increase the hunger and motivation to work for food, particularly in the HFD group. Each recording session consisted of a 5-minute baseline period, followed by structured trials (60 s inter-trial interval) for 60 minutes (Fig. 1, 60 trials) or 90 minutes (Fig. 2, 90 trials). An audible cue (tone, 2 kHz) was presented on each trial, at which point the mouse had 1 second to lick to trigger the lickspout for Ensure delivery via the lickspout. If no lick trigger (no tongue pressure exerted on the spout) occurred during the cue window, the mouse would not get any Ensure and must wait till the next audible cue (1 min later) to try again. An IR beam was used to measure the speed of free locomotion on the circular treadmill.

### Histology

Mice were terminally anesthetized using tribromoethanol (i.p. 250 mg/kg) and transcardially perfused with PBS and then formalin (10%). The tissue was fixed in 20% sucrose in PBS before extraction and microtome slicing (40 µm). Slides were stained with antifade mounting medium with DAPI (VECTASHIELD® HardSet™) before capturing the images with an Olympus VS120 slide scanner microscope.

### Analysis of head-fixed feeding assays

Data analysis was performed as described previously using custom software written in MATLAB (MathWorks) and Python (https://github.com/xzhang03/Photometry_analysis)^5,23^. Photometry signals in the photodetector trace were identified from LED pulses associated with illumination of the 465 nm LED and the median of the respective datapoints were extracted. The resulting trace (50 Hz sampling) was further filtered with a 10 Hz low-pass filter. The data were split into 60-s long traces beginning 10 s prior to the cue. We calculated baseline fluorescence (mean of 10 s pre-cue, denoted Fo) and fractional changes in fluorescence from the baseline (ΔF/Fo). Given that animals could voluntarily trigger the Ensure release repeatedly, with the number of total triggered trials depending on their hunger state and motivation, we considered only those recordings where at least 30 trials were triggered, unless otherwise stated, and for the analysis, we only selected the triggered trials for each session. The recordings from each animal from different days were concatenated to determine a single Z-score per animal, which was then used to determine the Z-score across the recordings.

Heatmaps were calculated by averaging across all usable trials across sessions from each mouse, and then a mean heatmap was computed across mice. Responses to Ensure during food consumption were estimated as the mean response during the peri-licking phase, 1-10 s post tone. For some analyses, as indicated in the Figure Legends, mean responses from 1-20 s post tone were considered. For the early meal analyses, we calculated the mean of the first 15 triggered trials, whereas the late meal was defined using the last 15 triggered trials. Lick rate during the cue was defined as the number of licks per second during the tone presentation. The lick rate during the reward window was calculated as the mean lick rate from 1-10s post tone. ΔZ-score (Fig. 2c, d, 3j) was calculated as a difference between the mean response during the peri-licking phase (1 – 10 s post tone) of the last 5 triggered trials and the first 5 triggered trials.

### Statistical analysis

Statistical analyses were performed in Python using the scipy.stats package. The numbers of recordings, animals, and statistics used are indicated in the figure legends. In summary, the statistical significance was determined with 1) two-way ANOVA and multiple comparisons performed using the Tukey HSD post hoc test in Fig 3h, k,, Supplementary Fig. 1h; 2) paired, two-tailed t-test between early and late meal trials Fig 1j Supplementary Fig. 1j, 3d; 3) unpaired, two-tailed Student’s t-test in Fig. 2b – d; Supplementary Fig. 1e, f; 4) one-way ANOVA with Tukey HSD post hoc test in Figure 3j; 5) linear mixed effects (LME) models (mixedlm of statsmodels Python module) to account for dependencies originating from repeated recording sessions of individual animals in Fig. 1i, 3i.

We fit a linear mixed-effects model to analyze the effects of diet and meal phase on the measured values while accounting for repeated measures (i.e., repeated sessions) within individual mice. The model included diet as a categorical fixed effect with “NCD” as the reference level and meal phase as an additional fixed effect. An interaction term between diet and meal phase was also included to assess whether the relationship between these factors influenced the outcome variable. To account for variability across individual mice, we specified “Mouse” as a random effect. The model was implemented using the formula: Value∼Diet×Meal phase+(1∣Mouse), where “Value” represents the dependent variable, “Diet” (NCD or HFD) and “Meal phase” (Early meal or Late meal) are fixed effects, and “Mouse” is the grouping factor for random intercepts. This approach allowed us to assess both the main effects and their interaction while controlling for inter-mouse variability. We used this model to analyze data presented in Figure 1i.

For diet-switched experiments, we fit a linear mixed-effects model to analyze the effects of Condition (a combination of start diet and second diet) and Meal Type (Early meal vs. Late meal) on the measured values while accounting for repeated measures within individual mice. The model included Condition as a categorical fixed effect with ‘NCD to NCD’ as the reference level, and Meal phase as an additional fixed effect. An interaction term between Condition and Meal phase was also included to assess whether the relationship between these factors influenced the outcome variable. To account for variability across individual mice, we specified “Mouse” as a random effect. The model was implemented using the formula: Value∼Condition×Meal phase+(1∣Mouse). Where “Value” represents the dependent variable, “Condition” (NCD -> NCD, HFD -> NCD, or HFD -> HFD) and “Meal phase” (Early meal or Late meal) are fixed effects, and “Mouse” serves as the grouping factor for random intercepts. This approach allowed us to assess both the main effects of Condition and Meal phase, as well as their interaction, while controlling for inter-mouse variability. The results from this model are presented in Figure 3i. Data representation with a scatter dot plot graph includes values of individual data points with mean values as a bar and standard error of the mean (s.e.m.) as error bars. Line plot graphs indicate the mean and show s.e.m. as a shaded region.

Significance was measured against an alpha value of 0.05 unless otherwise stated. *p < 0.05, **p < 0.01, ***p< 0.001, ****p<0.0001.

## Acknowledgments

This research was supported by a Walter Benjamin Fellowship from Deutsches Forschungsgemeinschaft PO 2925/1-1 (M.P), a Lefler Fellowship, a Charles A. King Trust Fellowship and NIH K99 DK134853 (S.X.Z.); R01 DK109930, DP1 AT010971, DP1 DK139958, a McKnight Scholar Award, and grants from the Boston Nutrition and Obesity Research Center (P30 DK046200), the Klarman Family Foundation, the Pew Innovation Fund and a Charles Robert Broderick III Phytocannabinoid Research Grant (M.L.A.). We thank all members of the Lehtinen and Andermann labs for fruitful discussions. We thank M. F. Hammell, J. Edelhaus, and L. Meschisen, for helping with animal care and histology, P. S. Sunkavalli, Z. A. Stolberg for behavioral experiments, H. Kucukdereli for technical support, P. Soden, P. Kalugin for further lab support, J. DeBolt, J. Shapiro, A. Pinilla, M. K. Lehtinen for administrative support.

## Author contributions

MP and SXZ designed the experiments with help from MLA. MP, SXZ, and MLA wrote the manuscript. MP conducted all surgeries. MP, JB, and CA performed the behavioral training and fiber photometry recordings. JB and CA performed all histology experiments. MP performed data analysis with guidance from SXZ and MLA.

---

**Supplementary Figure 1.**
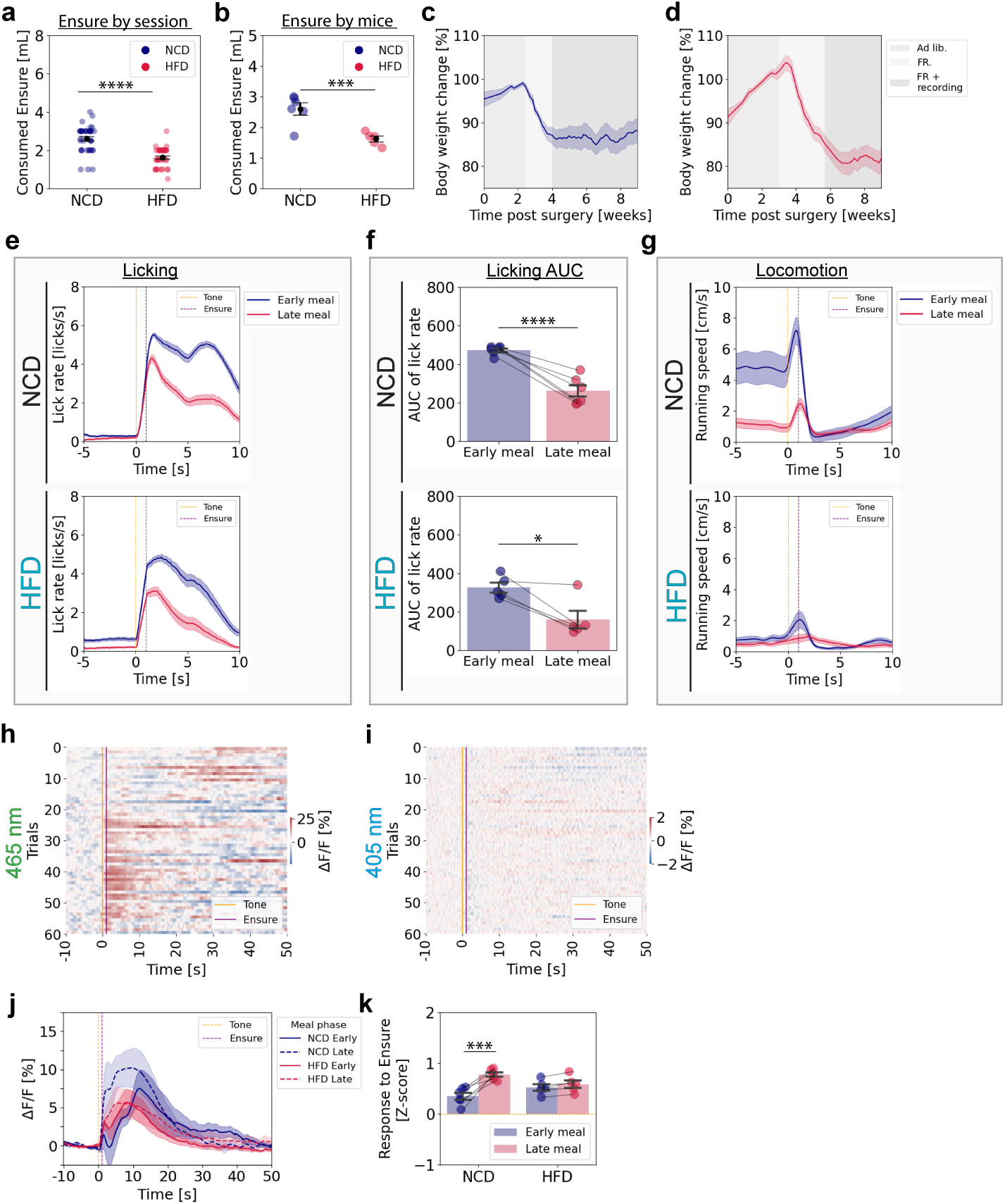
Validation of GCaMP6s photometry signals and behavioral dynamics during food restriction in NCD- and HFD-fed mice. **a, b,** Total Ensure consumption during the individual photometry recording sessions (**a**) and across individual mice (**b**) for NCD-fed and HFD-fed mice. **c, d**, Post-surgical bodyweight dynamics during ad libitum feeding (light gray), food restriction (FR, no shading), and continued food restriction maintenance during the photometry recording (FR + recording, dark shading) of NCD-fed (**c**) and HFD-fed (**d**) mice. **e**, Mean lick rate during early and late meal phases for NCD-fed (top panel) and HFD-fed (bottom panel) mice. **f**, Area under the curve (AUC) quantification of post-cue lick-rate, normalized to baseline pre-tone lick rate, for early and late meal phases of individual NCD-fed (top panel) and HFD-fed (bottom panel) mice. **g**, Mean running during the early and late meal phase of NCD-fed (top panel) and HFD-fed (bottom panel) mice. **h, i,** Representative heatmap of normalized ΔF/F GCaMP6s fluorescent responses detected using a 465 nm LED (**h**) and a 405 nm LED, reflecting movement artifacts and auto-fluorescence (**i**). Responses in each trial are normalized to the 10 s baseline before the cue onset. Note the range in **i** is smaller than in **a**. **j**, Early and late meal responses to Ensure for NCD-fed and HFD-fed animals, expressed as ΔF/F. **k,** Z-scored PVH^MC4R^ neuronal responses of each animal during early and late meal phases for all recordings in NCD-fed and HFD-fed mice, averaged from 1 – 20 s post-cue presentation. NCD: 6 mice; HFD: 5 mice. **a, b, f,** – unpaired, two-tailed Student t-test. **k,** Two-way Anova with Tukey post hoc test, for Early vs. Late meal within each condition and early phases between conditions. **a – g, j – k**, Data are represented as the mean ± s.e.m either as error bars (**a, b, f, k**) or as shaded area (**c, d, e, g, j**). **a, b, k**, * - P <0.05, *** - P <0.001, **** - P <0.0001.

**Supplementary Figure 2.**
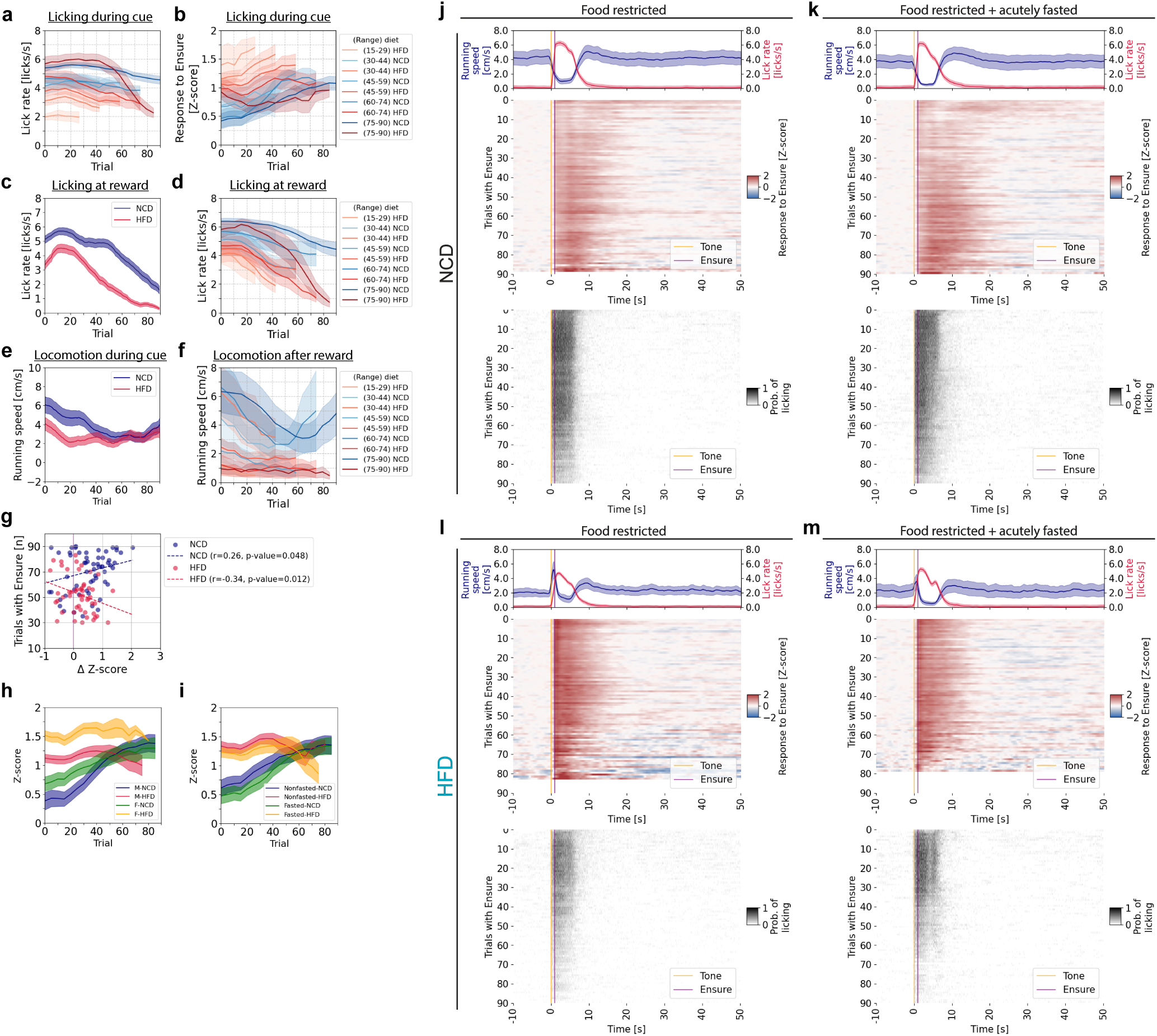
Diet- and fasting-dependent modulation of PVH^MC4R^ neuronal activity, licking behavior, and trial engagement in NCD- and HFD-fed mice. **a**, Lick rate during the 1 s cue window grouped by the number of triggered trials (ranges: 15–29, 30–44, 45–59, 60–74, 75–90) in NCD-fed (blue shade) and HFD-fed (red shade) mice. **b,** Single trial mean neuronal responses 1-9 s post cue (peri-licking) grouped by the number of triggered trials (ranges: 15–29, 30–44, 45–59, 60–74, 75–90) in NCD-fed (blue shade) and HFD-fed (red shade) mice. **c**, Single trial mean licking rate during the reward period (1 – 9 s post-cue) across 90 recorded trials from NCD-fed and HFD-fed groups (minimum 30 successfully triggered trials). **d**, Single trial mean licking rate during the reward (1 – 9 s post cue) across 90 recorded trails grouped by the number of triggered trials (ranges: 15–29, 30–44, 45–59, 60–74, 75–90) for NCD-fed (blue shading) and HFD-fed (red shading) mice. **e**, Single trial mean running speed across 90 recorded trails from NCD-fed and HFD-fed mice (minimum 30 successfully triggered trials). **f,** Single trial mean running speed during the pre-cue period across 90 recorded trails, grouped by the number of triggered trials (ranges: 15–29, 30–44, 45–59, 60–74, 75–90) for NCD-fed (blue shade) and HFD-fed (red shade) mice. **g**, Number of triggered trials out of the 90 possible trials recording session versus ΔZ-score between computed by subtracting the mean early meal response from the mean late meal response. The dashed line represents the linear relationship between meal size (triggered trials with Ensure) and the ΔZ-score (i.e. the change in response to Ensure from early to late in the meal). **h**, Single trial mean peri-licking PVH^MC4R^ responses (1 – 9 s post-cue) across 90 recorded trials from female (F) and male (M) NCD-fed and HFD-fed groups. Only recordings with at least 30 successfully triggered trials were included. M-NCD: 26 recordings / 3 mice; M-HFD: 31 recordings / 4 mice; F-NCD: 33 recordings / 4 mice; F-HFD: 23 recordings / 3 mice. **i**, Single-trial mean peri-licking PVH^MC4R^ responses (1 – 9 s post-cue) across 90 recorded trials for food restricted (nonfasted) and food restricted plus acute fasting (fasted) NCD-fed and HFD-fed groups. Only recordings with at least 30 successfully triggered trials were included. **j – m**, Heatmaps summarizing GCaMP6s photometry signals (top panel) and licking behavior (bottom panel) from PVH^MC4R^ neurons in food-restricted-only session (**j,l**) and in food restricted-plus acutely-fasted sessions (**k, m**) for NCD-fed (**j, k**) and HFD-fed (**l, m**) animals. Mean running speed and licking rates across all triggered trials are shown above the heatmaps. Only recordings with a minimum of 30 successfully triggered trials were included. Trial structure: 10 s baseline before cue (tone) onset (t = 0 s), followed by Ensure delivery at t = 1 s. **a – e**, Nonfasted NCD: 31 recordings / 7 mice (**i, j**); Food restricted plus acutely fasted NCD: 28 recordings / 7 mice (**i, k**); Nonfasted HFD: 31 recordings / 7 mice (**i, l**); Food restricted plus acutely fasted HFD: 23 recordings / 7 mice (**i, m**). **e – m**, Data are represented as the mean ± s.e.m. * - P <0.05, *** - P <0.001, **** - P <0.0001. **c, e, g,** NCD: 59 recordings / 7 mice; HFD: 54 recordings / 7 mice. **a, b, d, f,** Recordings for each meal size range: NCD (15–29): 0; HFD (15–29): 6; NCD (30–44): 6; HFD (30–44): 13; NCD (45–59): 10; HFD (45–59): 23; NCD (60–74): 13; HFD (60–74): 7; NCD (75–90): 28; HFD (75–90): 4.

**Supplementary Figure 3.**
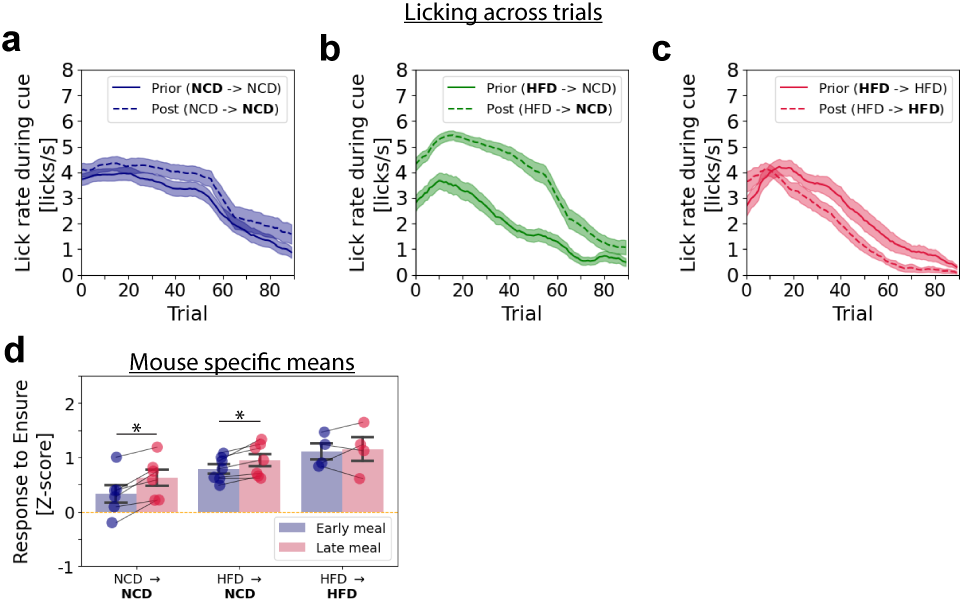
Impact of diet switch on cue-evoked licking behavior and PVH^MC4R^ dynamics during early and late meal phases. **a – c**, Single-trial mean lick rate during cue presentation (0 – 1 s post-cue onset) across 90 trials averaged across recordings with at least 30 successfully triggered trials in the NCD to NCD (**a**), HFD to NCD (**b**), and HFD to HFD (**c**) groups, comparing pre-diet-switch (solid line) and post-diet-switch (dashed line) conditions. **a – c,** Recording and animal numbers during exposure to the first diet: **NCD** to NCD: 44 recordings / 6 mice; **HFD** to NCD: 52 recordings / 7 mice; **HFD** to HFD: 35 recordings / 4 mice. Recording and animal numbers during exposure to the second diet: NCD to **NCD**: 43 recordings / 6 mice; HFD to **NCD**: 42 recordings / 7 mice; HFD to **HFD**: 23 recordings / 4 mice. **d**, Mean Z-scored PVH^MC4R^ neuronal responses during early (first 15 trials) and late meal (last 15 trials) phases of the meal, for individual mice (averaged across sessions per mouse) across the following conditions post-diet switch: NCD to NCD, HFD to NCD, HFD to HFD. Paired, two-tailed t-test between early-meal and late-meal trials. **a – d**, Data are represented as the mean ± s.e.m across sessions (**a – c**) or across mice (**d**). * - P <0.05, ** - P <0.01, *** - P <0.001.

## References

1. Andermann, M. L. & Lowell, B. B. Toward a Wiring Diagram Understanding of Appetite Control. Neuron 95, 757–778 (2017).

2. Garfield, A. S. et al. A neural basis for melanocortin-4 receptor–regulated appetite. Nat. Neurosci. 18, 863–871 (2015).

3. Sweeney, P., Gimenez, L. E., Hernandez, C. C. & Cone, R. D. Targeting the central melanocortin system for the treatment of metabolic disorders. Nat. Rev. Endocrinol. 19, 507–519 (2023).

4. Balthasar, N. et al. Divergence of melanocortin pathways in the control of food intake and energy expenditure. Cell 123, 493–505 (2005).

5. Zhang, S. X. et al. Stochastic neuropeptide signals compete to calibrate the rate of satiation. Nature 637, 137–144 (2025).

6. De Solis, A. J. et al. Reciprocal activity of AgRP and POMC neurons governs coordinated control of feeding and metabolism. Nat. Metab. 6, 473–493 (2024).

7. Krashes, M. J. et al. Rapid, reversible activation of AgRP neurons drives feeding behavior in mice. J. Clin. Invest. 121, 1424–1428 (2011).

8. van den Pol, A. N. Neuropeptide Transmission in Brain Circuits. Neuron 76, 98–115 (2012).

9. Steculorum, S. M. et al. AgRP Neurons Control Systemic Insulin Sensitivity via Myostatin Expression in Brown Adipose Tissue. Cell 165, 125–138 (2016).

10. Engström Ruud, L., Pereira, M. M. A., de Solis, A. J., Fenselau, H. & Brüning, J. C. NPY mediates the rapid feeding and glucose metabolism regulatory functions of AgRP neurons. Nat. Commun. 11, 442 (2020).

11. Tong, Q., Ye, C.-P., Jones, J. E., Elmquist, J. K. & Lowell, B. B. Synaptic release of GABA by AgRP neurons is required for normal regulation of energy balance. Nat. Neurosci. 11, 998–1000 (2008).

12. Wu, Q., Boyle, M. P. & Palmiter, R. D. Loss of GABAergic signaling by AgRP neurons to the parabrachial nucleus leads to starvation. Cell 137, 1225–1234 (2009).

13. Gropp, E. et al. Agouti-related peptide–expressing neurons are mandatory for feeding. Nat. Neurosci. 8, 1289–1291 (2005).

14. Luquet, S., Perez, F. A., Hnasko, T. S. & Palmiter, R. D. NPY/AgRP neurons are essential for feeding in adult mice but can be ablated in neonates. Science 310, 683–685 (2005).

15. Mountjoy, K. G., Robbins, L. S., Mortrud, M. T. & Cone, R. D. The Cloning of a Family of Genes That Encode the Melanocortin Receptors. Science 257, 1248–1251 (1992).

16. Brüning, J. C. & Fenselau, H. Integrative neurocircuits that control metabolism and food intake. Science 381, eabl7398 (2023).

17. Friedman, J. M. Leptin and the endocrine control of energy balance. Nat. Metab. 1, 754– 764 (2019).

18. Ruud, J., Steculorum, S. M. & Brüning, J. C. Neuronal control of peripheral insulin sensitivity and glucose metabolism. Nat. Commun. 8, 15259 (2017).

19. Cowley, M. A. et al. The Distribution and Mechanism of Action of Ghrelin in the CNS Demonstrates a Novel Hypothalamic Circuit Regulating Energy Homeostasis. Neuron 37, 649–661 (2003).

20. D’Agostino, G. & Diano, S. alpha-Melanocyte stimulating hormone: production and degradation. J. Mol. Med. 88, 1195–1201 (2010).

21. Xu, S. et al. Behavioral state coding by molecularly defined paraventricular hypothalamic cell type ensembles. Science 370, (2020).

22. Molden, B. M., Cooney, K. A., West, K., Van Der Ploeg, L. H. T. & Baldini, G. Temporal cAMP Signaling Selectivity by Natural and Synthetic MC4R Agonists. Mol. Endocrinol. 29, 1619–1633 (2015).

23. Zhang, G.-W. et al. Medial Preoptic Area Antagonistically Mediates Stress-induced Anxiety and Parental Behavior. Nat. Neurosci. 24, 516–528 (2021).

24. Jais, A. & Brüning, J. C. Arcuate Nucleus-Dependent Regulation of Metabolism-Pathways to Obesity and Diabetes Mellitus. Endocr. Rev. 43, 314–328 (2022).

25. Beutler, L. R. et al. Obesity causes selective and long-lasting desensitization of AgRP neurons to dietary fat. eLife 9,.

26. Loos, R. J. F. & Yeo, G. S. H. The genetics of obesity: from discovery to biology. Nat. Rev. Genet. 23, 120–133 (2022).

27. Huszar, D. et al. Targeted disruption of the melanocortin-4 receptor results in obesity in mice. Cell 88, 131–141 (1997).

28. Krude, H. et al. Severe early-onset obesity, adrenal insufficiency and red hair pigmentation caused by POMC mutations in humans. Nat. Genet. 19, 155–157 (1998).

29. Yeo, G. S. H. et al. A frameshift mutation in MC4R associated with dominantly inherited human obesity. Nat. Genet. 20, 111–112 (1998).

30. Vaisse, C., Clement, K., Guy-Grand, B. & Froguel, P. A frameshift mutation in human MC4R is associated with a dominant form of obesity. Nat. Genet. 20, 113–114 (1998).

31. Jackson, R. S. et al. Obesity and impaired prohormone processing associated with mutations in the human prohormone convertase 1 gene. Nat. Genet. 16, 303–306 (1997).

32. Montague, C. T. et al. Congenital leptin deficiency is associated with severe early-onset obesity in humans. Nature 387, 903–908 (1997).

33. Kühnen, P. et al. Proopiomelanocortin Deficiency Treated with a Melanocortin-4 Receptor Agonist. N. Engl. J. Med. 375, 240–246 (2016).

34. Ryan, D. H. Setmelanotide: what does it mean for clinical care of patients with obesity? Lancet Diabetes Endocrinol. 8, 933–935 (2020).

35. Farooqi, I. S. & O’Rahilly, S. Mutations in ligands and receptors of the leptin–melanocortin pathway that lead to obesity. Nat. Clin. Pract. Endocrinol. Metab. 4, 569– 577 (2008).

36. Kievit, P. et al. Chronic Treatment With a Melanocortin-4 Receptor Agonist Causes Weight Loss, Reduces Insulin Resistance, and Improves Cardiovascular Function in Diet-Induced Obese Rhesus Macaques. Diabetes 62, 490–497 (2013).

37. Li, H. et al. The melanocortin action is biased toward protection from weight loss in mice. Nat. Commun. 14, 2200 (2023).

38. Mazzone, C. M. et al. High-fat food biases hypothalamic and mesolimbic expression of consummatory drives. Nat. Neurosci. 23, 1253–1266 (2020).

39. Enriori, P. J. et al. Diet-Induced Obesity Causes Severe but Reversible Leptin Resistance in Arcuate Melanocortin Neurons. Cell Metab. 5, 181–194 (2007).

40. Briggs, D. I., Enriori, P. J., Lemus, M. B., Cowley, M. A. & Andrews, Z. B. Diet-Induced Obesity Causes Ghrelin Resistance in Arcuate NPY/AgRP Neurons. Endocrinology 151, 4745–4755 (2010).

41. Briggs, D. I. et al. Evidence that diet-induced hyperleptinemia, but not hypothalamic gliosis, causes ghrelin resistance in NPY/AgRP neurons of male mice. Endocrinology 155, 2411–2422 (2014).

42. Paeger, L. et al. Energy imbalance alters Ca2+ handling and excitability of POMC neurons. eLife 6, e25641 (2017).

43. Beutler, L. R. et al. Dynamics of Gut-Brain Communication Underlying Hunger. Neuron 96, 461–475.e5 (2017).

44. Chen, Y., Lin, Y.-C., Kuo, T.-W. & Knight, Z. A. Sensory Detection of Food Rapidly Modulates Arcuate Feeding Circuits. Cell 160, 829–841 (2015).

45. Chen, Y. et al. Sustained NPY signaling enables AgRP neurons to drive feeding. eLife 8, e46348 (2019).

46. Dodd, G. T. et al. Intranasal Targeting of Hypothalamic PTP1B and TCPTP Reinstates Leptin and Insulin Sensitivity and Promotes Weight Loss in Obesity. Cell Rep. 28, 2905–2922.e5 (2019).

47. Dodd, G. T. et al. Insulin signaling in AgRP neurons regulates meal size to limit glucose excursions and insulin resistance. Sci. Adv. 7, eabf4100 (2021).

48. Nyamugenda, E. et al. Selective Survival of Sim1/MC4R Neurons in Diet-Induced Obesity. iScience 23, 101114 (2020).

49. Cooney, K. A., Molden, B. M., Kowalczyk, N. S., Russell, S. & Baldini, G. Lipid stress inhibits endocytosis of melanocortin-4 receptor from modified clathrin-enriched sites and impairs receptor desensitization. J. Biol. Chem. 292, 17731–17745 (2017).

50. Briggs, D. I. et al. Calorie-Restricted Weight Loss Reverses High-Fat Diet-Induced Ghrelin Resistance, Which Contributes to Rebound Weight Gain in a Ghrelin-Dependent Manner. Endocrinology 154, 709–717 (2013).

51. Baver, S. B. et al. Leptin Modulates the Intrinsic Excitability of AgRP/NPY Neurons in the Arcuate Nucleus of the Hypothalamus. J. Neurosci. 34, 5486–5496 (2014).

52. Ottaway, N. et al. Diet-induced obese mice retain endogenous leptin action. Cell Metab. 21, 877–882 (2015).

53. Zhao, S. et al. Partial Leptin Reduction as an Insulin Sensitization and Weight Loss Strategy. Cell Metab. 30, 706–719.e6 (2019).

54. Berthoud, H.-R., Münzberg, H. & Morrison, C. D. Blaming the Brain for Obesity: Integration of Hedonic and Homeostatic Mechanisms. Gastroenterology 152, 1728– 1738 (2017).

